# Evaluating live microbiota biobanking using an *ex vivo* microbiome assay and metaproteomics

**DOI:** 10.1101/2021.09.30.462621

**Authors:** Xu Zhang, Krystal Walker, Janice Mayne, Leyuan Li, Zhibin Ning, Alain Stintzi, Daniel Figeys

## Abstract

Biobanking of live microbiota is becoming indispensable for mechanistic and clinical investigations of drug-microbiome interactions and fecal microbiota transplantation. However, there is a lack of methods to rapidly and systematically evaluate whether the biobanked microbiota maintains their cultivability and functional activity. In this study, we use a rapid *ex vivo* microbiome assay and metaproteomics to evaluate the cultivability and the functional responses of biobanked microbiota to treatment with a prebiotic (fructo-oligosaccharide, FOS). Our results indicate that the microbiota cultivability and their functional responses to FOS treatment were well maintained by freezing in a deoxygenated glycerol buffer at -80°C for 12 months. We also demonstrate that the fecal microbiota is functionally stable for 48 hours on ice in a deoxygenated glycerol buffer, allowing off-site fecal sample collection and shipping to laboratory for live microbiota biobanking. This study provides a method for rapid evaluation of the cultivability of biobanked live microbiota. Our results show minimal detrimental influences of long-term freezing in deoxygenated glycerol buffer on the cultivability of fecal microbiota.

## Introduction

Increasing evidence associates the gut microbiota with human health and the development of various diseases, including both intestinal and non-intestinal disorders ^1-3^. Fecal microbiota transplantation (FMT) has been used to treat recurrent *Clostridium difficile* infection (CDI), inflammatory bowel diseases (IBD) ^4, 5^ and to overcome resistance to anti-PD1 therapy in melanoma patients ^6, 7^. Small molecules, prebiotics and dietary components are also being studied in high-throughput *in vitro* microbiome screens to identify potential therapeutics targeting the gut microbiota ^8-12^ and to study the effects of the microbiome on these compounds ^13^. Although some successes have emerged in therapeutic interventions of gut microbiota for disease management ^14-16^, there is a need for systematic and robust study of microbiome-therapeutic interactions. Biobanked live microbiota are increasingly used for FMT and in high-throughput screening for new therapeutics, but most studies do not investigate whether biobanked microbiota maintains their functionalities.

Biobanked stools stored in the presence of preservation chemicals, such as glycerol, are commonly used for either microbiome assay, fermentation or FMT applications ^13, 17, 18^. Using the bacterial cell counting approach, Guerin-Danan *et al*. demonstrated that freezing in glycerol buffer had no effect on aerobes in fecal samples, while decreased the survival of anaerobes, but did not exceed the inter-individual variations ^19^. In a mouse FMT experiment, Ericsson *et al*. demonstrated that FMT using frozen feces performed similarly to fresh feces in maintaining the richness, diversity, and composition of donor microbiota ^20^. Utilizing a propidium monoazide (PMA)-based live bacteria-specific P18CR and sequencing approach, Papanicolas *et al*. showed that freeze-thaw reduced the microbial viability to ∼20% in feces, although the composition of viable microbiota was not significantly altered ^21^. In addition, the exposure of fecal samples to ambient air further reduced viability and could lead to >10-fold reductions in the abundance of important commensal bacteria, such as *Faecalibacterium prausnitzii* ^21^. Therefore, biobanking of live microbiota needs to be systematically evaluated to determine whether the viability and functional activity of microbiota are maintained.

High-throughput whole-microbiome culture methods have been recently developed ^11, 13^. For example, we reported a rapid assay for individual microbiome (RapidAIM) ^11^, which maintains the functional profiles of individual gut microbiomes. RapidAIM has been used to screen, evaluate, and reclassify the modulating effects of drugs, natural compounds, and dietary components against individual human microbiomes ^8, 9^. In this study, we used the RapidAIM assay and quantitative metaproteomics to determine whether freezing and delayed sample processing of human gut microbiota affect their cultivability and functional responses to treatments. Briefly, fresh and frozen microbiomes isolated from human stools were cultured in the RapidAIM assay with or without fructo-oligosaccharides (FOS), a prebiotic with known effects on microbiota ^22-24^. We showed that microbiome samples stored in deoxygenated, buffered 10% glycerol at -80°C were stable and maintained their functional responses to FOS treatment for up to 1 year. In addition, we demonstrated that gut microbiomes were stable on ice with glycerol-based preservation buffer for 48 hours prior to sample processing and biobanking. This study provides a convenient and rapid approach for evaluating live microbiota and reveals that freezing in glycerol-based preservation buffer at -80°C minimally affects the cultivability and activity of gut microbiomes.

## Results

### Freezing does not change the functional individuality of cultured gut microbiome

We first evaluated whether frozen biobanked fecal microbiota maintain their cultivability and functions. Briefly, fresh stools from three adult volunteers were collected, processed to make a final concentration of 20% (w/v) fecal slurry in 10% (v/v) glycerol buffer (containing 1 mg/ml L-cysteine and pre-equilibrated in anaerobic workstation over night). The fecal slurry was aliquoted and stored at -80°C up to 1 year with testing at 0, 1, 2, 4, 8, 16, 24, and 52 weeks (1 year) by RapidAIM (Figure 1a). RapidAIM culturing was performed with or without 5 mg/ml of FOS for 24 hours in an anaerobic workstation. The microbiome cultures as well as the uncultured baseline samples were analysed by shotgun metaproteomic to examine the functional microbiome profiles overtime.

**Figure 1.**
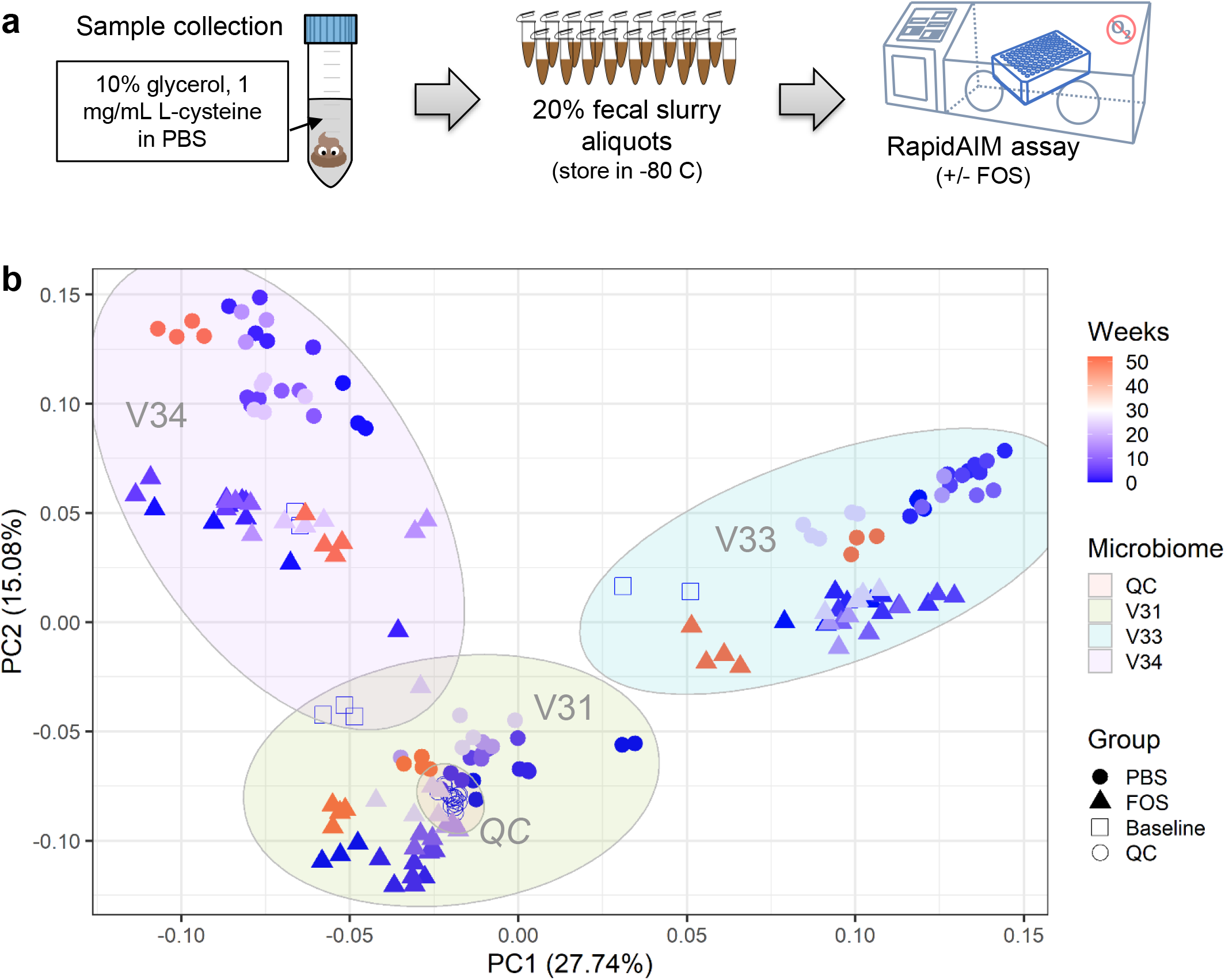
Evaluation of the cultivability of frozen biobanked fecal microbiota. (a) experimental workflow; (b) PCA score plot of quantified protein groups. The log10-tranformed LFQ intensities of protein groups that were quantified in >50% samples were used for PCA analysis.

In this study, 181 MS raw files were generated, including 16 quality control (QC) MS runs. Protein identification with MetaLab software yielded 83,273 unique peptide sequences and 20,304 protein groups, with an average MS spectra identification rate of 42.1 ± 6.9% (mean ± standard deviation). To examine the overall protein expression profiles, protein groups that were quantified in at least 50% of the samples were used for principal component analysis (PCA, Figure 1b). As shown in the PCA score plot, QC runs clustered closely together, indicating high quality and consistency of the metaproteomic data throughout the year (Figure 1b). Overall, the samples segregated into three clusters according to the individual origin of microbiomes tested. Both pre- and post-cultured microbiome samples of the same individual clustered together, suggesting adequate maintenance of the microbiome functional profiles ^10^. Within each individual microbiome, the treatment of FOS led to obvious separation (as represented by the second principal component, PC2) (Figure 1b). These findings suggest that the individuality of microbiome functions was well maintained by biobanking, *ex vivo* culturing, as well as the treatment with prebiotic FOS.

### Frozen biobanked microbiota maintains the functional responses to FOS in RapidAIM

A total of 19,134 (94%) identified gut microbial protein groups were annotated with the Clusters of Orthologous Groups of proteins (COGs) database, representing 1214 COGs and 24 COG categories. The relative abundances of COGs were then used for PCA analysis, which again demonstrated consistency of individual microbiome functional responses to culturing, biobanking and treatment with FOS (Figure 2). Obvious separation of samples treated with and without FOS was observed along the second PC for all three microbiomes (Figure 2). We then calculated the abundance distributions at COG category level for each group, which showed highly similar patterns for all microbiomes at all different time points (Figure 3). Interestingly, consistent functional responses of cultured microbiomes to FOS could be observed for all tested time points, indicating well maintained cultivability and activity of the biobanked microbiomes.

**Figure 2.**
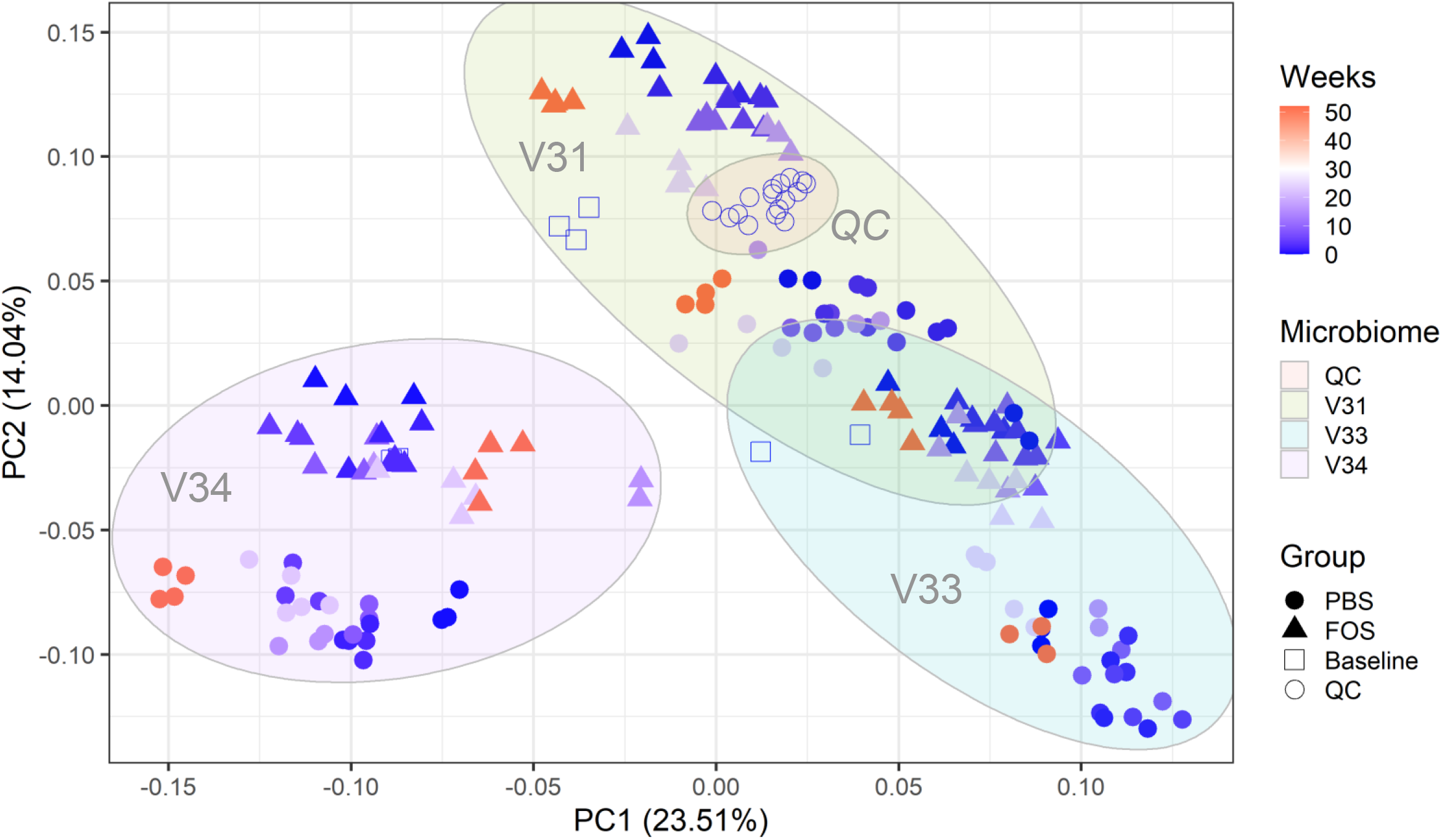
PCA score plot of quantified functional orthologs of baseline and cultured microbiomes using fresh or frozen biobanked stools. The log10-tranformed LFQ intensities of COGs that were quantified in >50% samples were used for PCA analysis.

**Figure 3.**
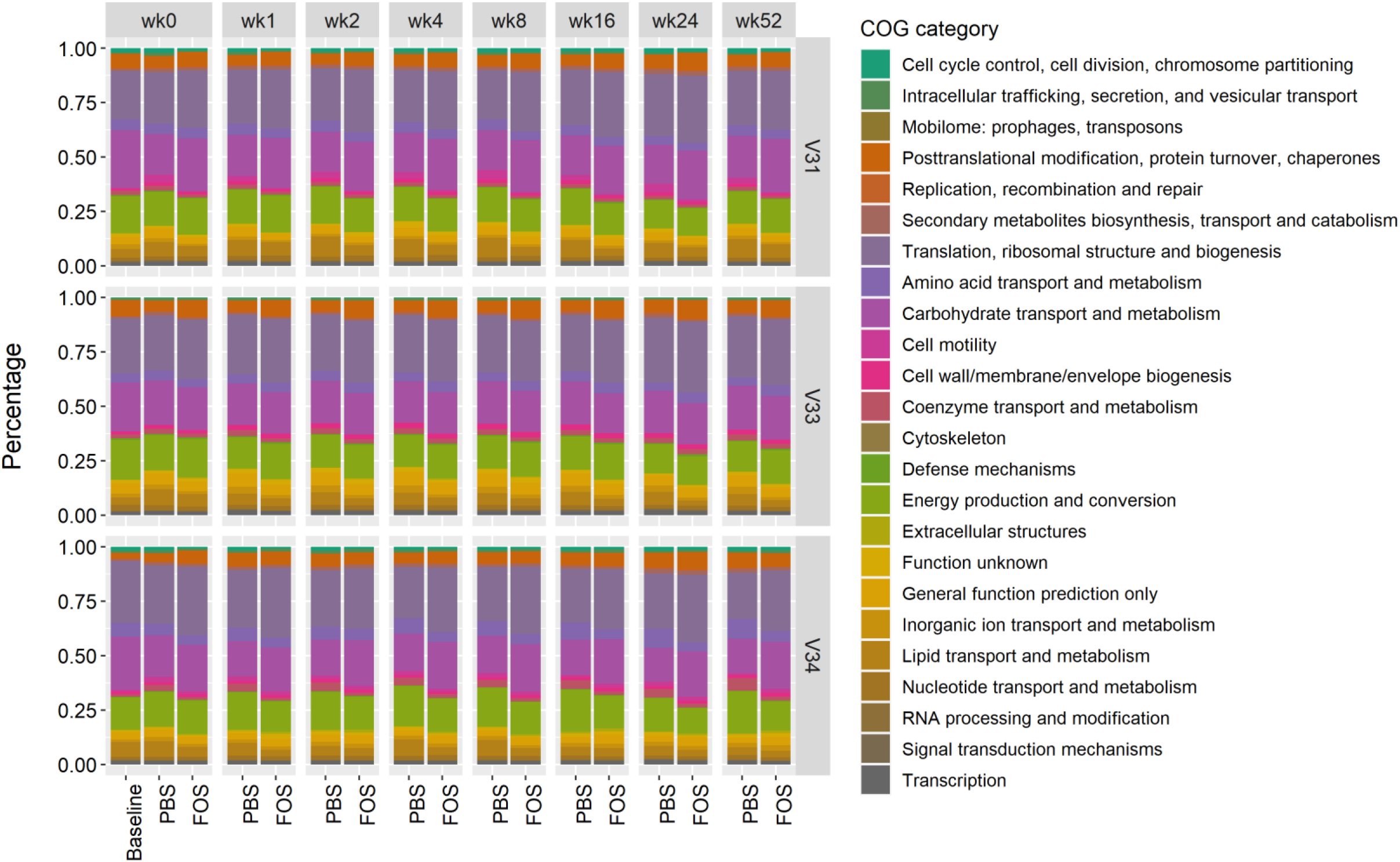
Abundance distribution of quantified COG categories in baseline and cultured microbiomes using fresh or frozen biobanked stools. Group average of LFQ intensities were used for plotting.

### Frozen biobanked microbiota maintains the taxonomic responses to FOS in RapidAIM

Quantitative taxonomic analysis was performed using identified distinctive peptides that were determined with a lowest common ancestor (LCA) approach. In total, 17 phyla from all 4 superkingdoms, 25 classes, 36 orders, 49 families, 74 genera, and 163 species were quantified using a threshold of ≥3 distinctive peptides. Overall phylum-level distribution of the pre- and post-cultured microbiomes showed high consistency of different groups and was distinct for different individuals (Figure 4). FOS is known to elevate the growth of Actinobacteria both *in vitro* and *in vivo* ^22-24^. Accordingly, consistent responses of the phylum Actinobacteria to FOS treatment were observed across all time points for all three tested microbiomes (Figure 5). Genus level analysis also showed that the two most abundant genera from Actinobacteria, *Collinsella* and *Bifidobacterium*, were consistently increased by FOS treatment (data not shown). The relative abundances of several phyla, such as Verrucomicrobia and Fusobacteria, were found to be responsive to freezing in certain microbiomes (Figure 4). In microbiome V31, the relative abundance of Verrucomicrobia was increased in the cultured samples using biobanked stools compared to those using fresh stools. Similarly, in microbiome V33, Fusobacteria were found to increase when frozen inoculums were used. These increases of Verrucomicrobia or Fusobacteria using frozen inoculums were recovered with the supplementation of FOS. These findings suggest that biobanking might lead to growth advantages for some bacterial species in the microbial community.

**Figure 4.**
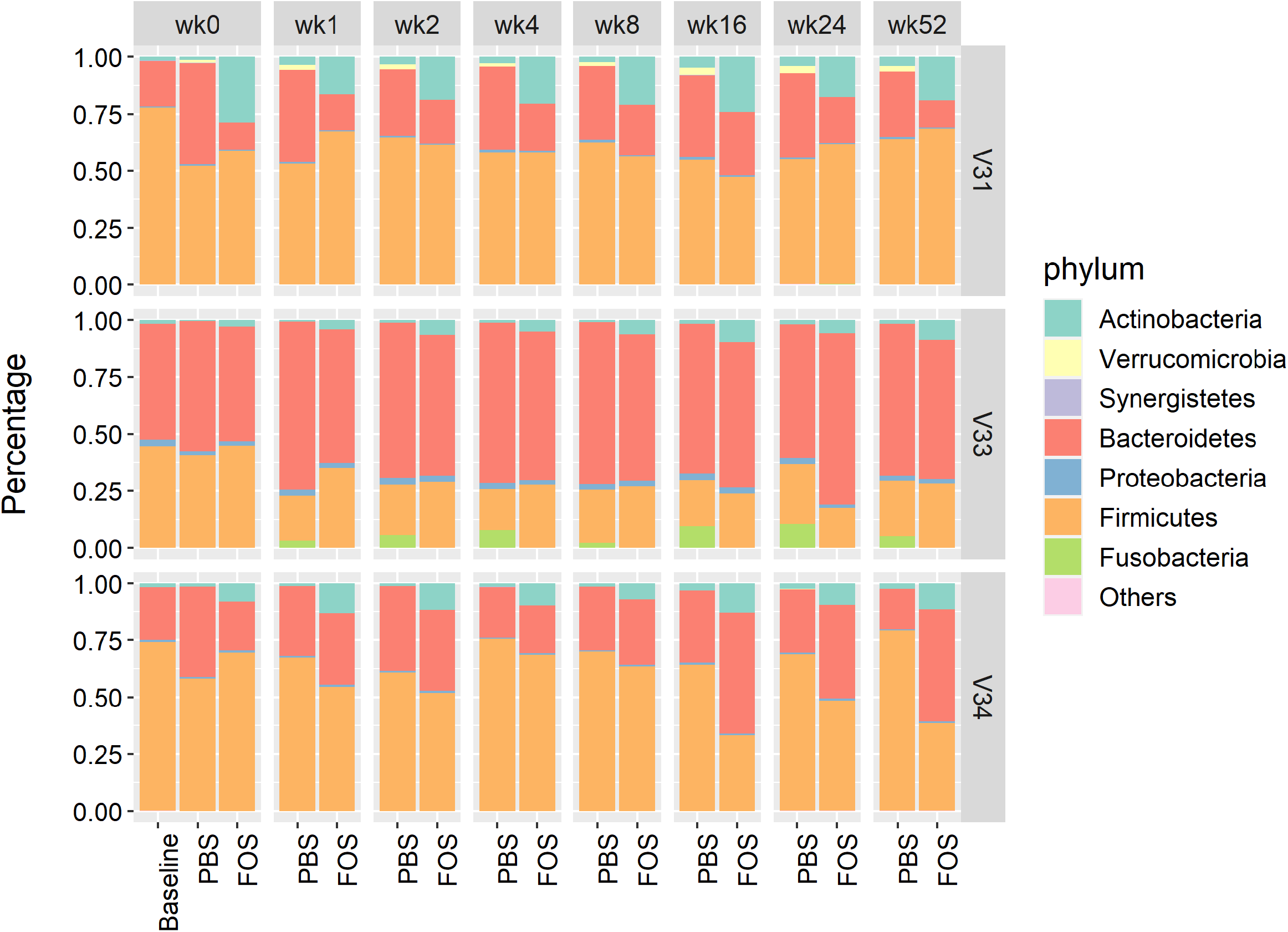
Phylum level composition of baseline and cultured microbiomes using fresh or frozen biobanked stools. Group average of relative abundances was used for plotting.

**Figure 5.**
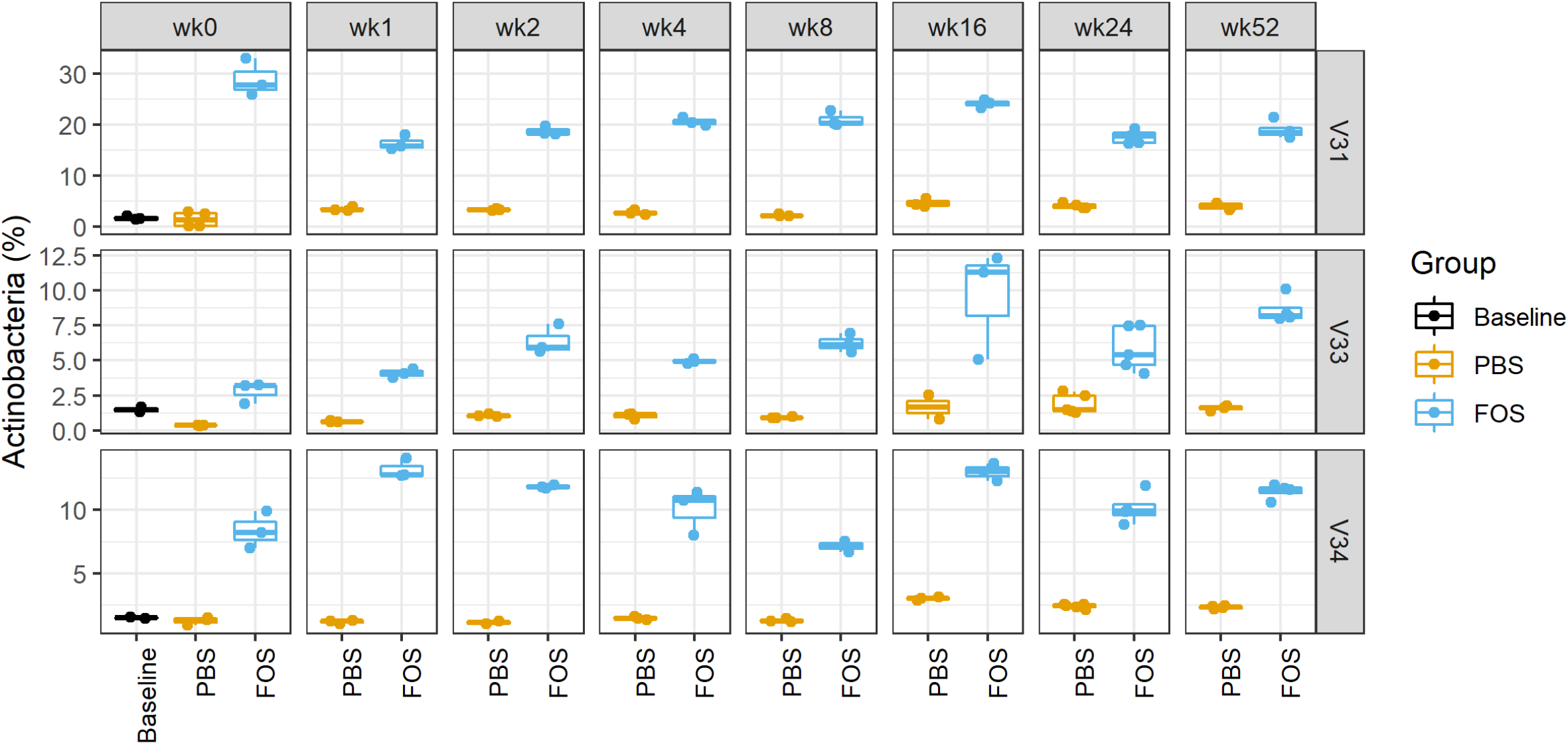
Relative abundances of Actinobacteria in baseline and cultured microbiomes using fresh or frozen biobanked stools. Box plots were generated using *ggplot2* with data points overlapped with the boxes. The first and third quantiles were indicated as the box width and median was also displayed in the middle of box. Upper and lower whiskers indicate the smallest value ≥1.5 interquartile range (IQR) and largest value ≤1.5 IQR, respectively.

### Microbiota cultivability is better maintained on ice with buffer than on dry ice

During biobanking, there is usually a delay prior to sample processing as the stools are usually collected off-site and shipped/transferred to the laboratory. We thereby evaluated whether fecal microbiota samples are stable during a delayed sample processing. We first compared stools that are (1) processed immediately, (2) stored on ice for 6 hours in deoxygenated glycerol buffer, and (3) directly stored on dry ice for 6 hours. Each of the stools was processed and cultured using RapidAIM assay with or without the treatment of FOS, and cultured samples were collected for metaproteomic analyses. PCA analysis of quantified protein groups showed that, for all the 3 tested microbiomes, most variations were found to be contributed by FOS treatment, however the samples from stools that were kept on dry ice were separated from fresh stools or stools that were kept on ice (Figure 6a). Taxonomic analysis also showed that the phylum level composition was more similar between fresh and ice-stored microbiomes than that between fresh and dry ice-stored microbiomes (Figure 6b). Interestingly, Fusobacteria were also found to be increased in V33 samples with dry ice-stored stools as inoculum, but not in those with ice-stored stools as inoculum (Figure 6b). These findings suggest that temporary storage of fecal samples on ice with deoxygenated glycerol buffer is better than on dry ice for maintaining the cultivability of microbiomes.

**Figure 6.**
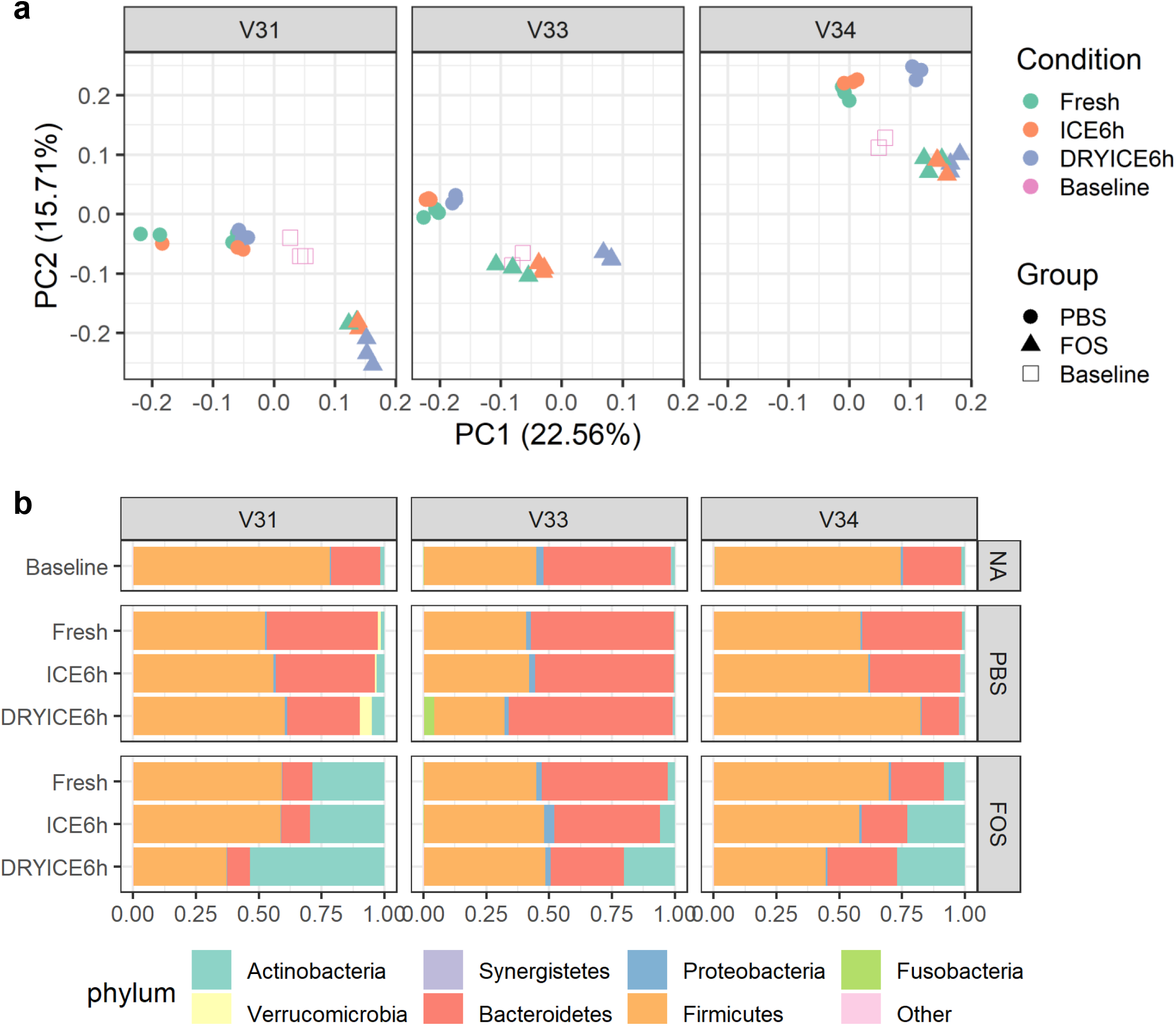
Effects of delayed fecal sample processing on the cultivability of microbiomes. (a) PCA score plots of quantified protein groups. (b) Phylum level taxonomic compositions as calculated using metaproteomic data. Group average of relative abundances was used for plotting.

We also evaluated whether fecal samples can be stored on ice for up to 48 hours mimicking a two-day shipping procedure for sample collection. Briefly, we simulated the shipping process by keeping stools in shipping packages with ice packs for 0, 24 and 48 hours, respectively, prior to processing and RapidAIM culturing. As shown in Figure 7a, while there was a shift along with the increase of shipping hours at PC2 (explained 7.26% of the total variations), most of the variations (57.12%) of the overall microbiome protein expressions were contributed by the treatment by FOS. Taxonomic analysis also showed that relatively stable phylum level composition of cultured microbiomes was obtained for up to 48 hours, and consistent responses of cultured microbiomes to FOS treatment were achieved (Figure 7b).

**Figure 7.**
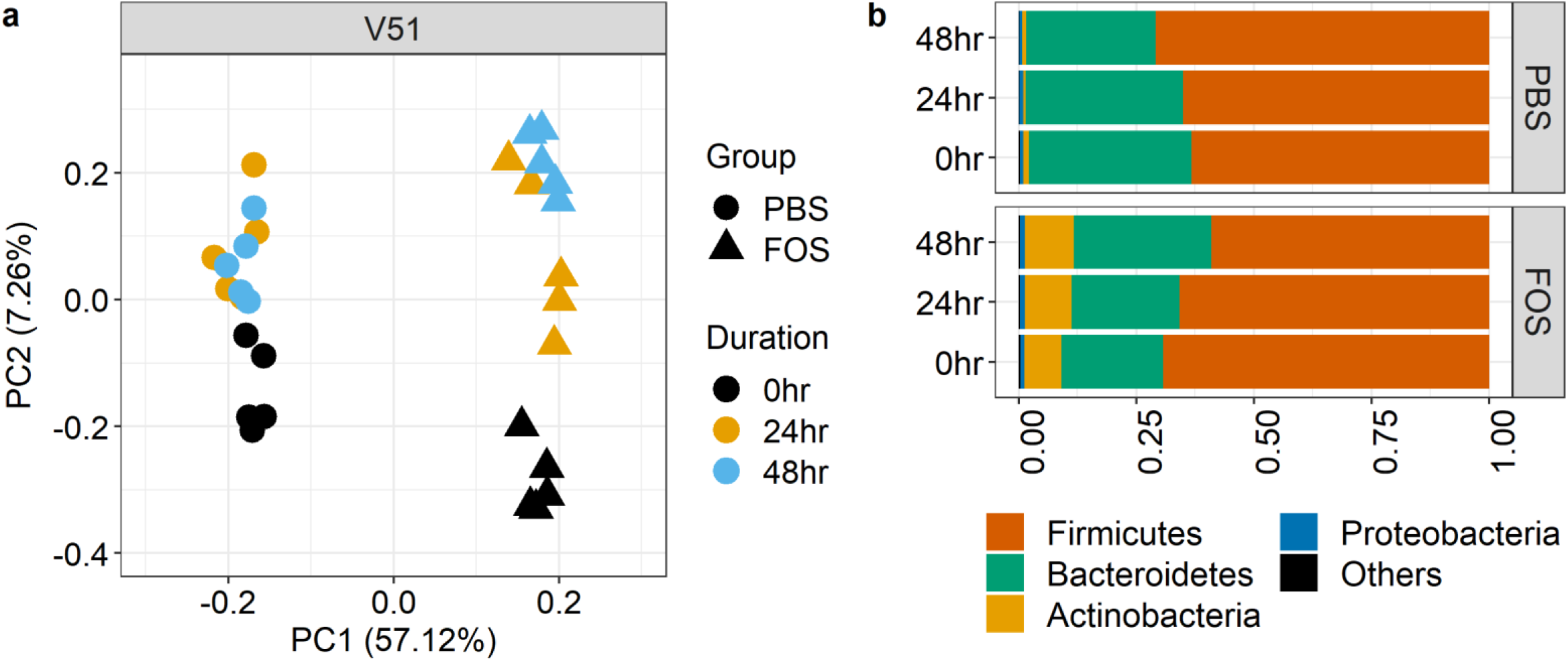
Influence of two-day delayed processing on the cultivability of microbiomes. (a) PCA score plots of quantified protein groups. (b) Phylum level taxonomic compositions as calculated using metaproteomic data. Group average of relative abundances was used for plotting.

## Discussion

The development of novel intervention approaches targeting the gut microbiome requires a high throughput screening of potential therapeutics against the microbiome, which leads to the increasing use of live microbiota ^25^. Due to the operational challenges for using fresh stools, most studies used frozen biobanked samples. However, evaluation of the functionality of frozen microbiotas has not been standardized and was missed in most studies. A few studies evaluated the performance of frozen microbiota using animal models and FMT ^20, 21^. However, animal experiments are expensive and time consuming. In this study, we used the RapidAIM assay ^11^ and metaproteomics to evaluate over one year whether frozen fecal microbiota in deoxygenated glycerol buffer maintains the microbiome cultivability and functional activity.

RapidAIM is an *in vitro* assay of individual microbiomes, which has been applied to study various compounds, such as FDA-approved drugs, resistant starches, and natural compounds, for their effects on human gut microbiomes ^8-11^. We have previously shown that RapidAIM maintains the functional and compositional profiles of microbiome and recapitulates known *in vivo* effects of drugs, such as metformin ^10^. Here, microbiomes cultured in the RapidAIM assay with and without a compound (i.e., FOS) with known microbiota-modulating effects were compared to baseline/uncultured microbiome samples (i.e., baseline, non-treated, and FOS-treated groups). We used quantitative metaproteomics to measure the expressed proteins and functions, because we were particularly interested in the functional state of the microbiomes, which is best assessed at the protein level instead of DNA level (16S rRNA gene sequencing or metagenomics) ^26^. Moreover, recent development of MS-based deep metaproteomics approaches ^26-28^ allows rapid and comprehensive profiling of the functions of the cultured microbiomes in this study. In agreement with previous FMT and animal studies ^20, 21^, we did not observe obvious detrimental effects on the functional profiles of the microbiomes frozen up to a year. Consistent responses of microbiome to FOS treatment, including the upregulation of Actinobacteria, were achieved using either fresh or frozen fecal samples.

Time spent in transporting and processing the stools is an important factor that may affect the viability of microbiota for biobanking. Gratton *et al*. reported that temperatures during transportation markedly affected the metabolite profiles of crude feces due to the microbial fermentation, which was reduced at 4°C ^29^. Fecal samples frozen at -20°C showed increased levels of amino acids, nicotinate and decreased levels of short-chain fatty acids (SCFAs) when compared to fresh samples ^29^. Therefore, Gheorghe *et al*. recommended that fecal samples should be kept at 4°C or on ice after collection and during transportation to better maintain the viability of microbiota for FMT ^30^. Burz *et al*. provided a comprehensive guideline in handling and storing stools for FMT through a systemic evaluation using 16S rRNA sequencing, metabolomic fingerprinting and flow cytometry, which also suggested that fresh samples should refrigerated at 4 °C if the transformation to transplant process exceeded 24 hours ^31^. Accordingly, in the current study, we demonstrated that the cultivability of microbiome and functional responses to treatment with FOS were better maintained in deoxygenated buffer on ice than on dry ice, and the microbiome was stable on ice in buffer for 48 hours. In contrast, cultured microbiomes using feces stored on dry ice showed marked changes in taxonomic compositions, even at phylum level, when compared to those using fresh feces. These findings confirm that fecal samples can be collected off-site, stored on ice, and shipped to laboratory using a standard two-day courier for further processing. Altogether, this study suggests the feasibility of using properly biobanked live microbiota for applications, such as drug-microbiome interaction study or FMT.

It is noteworthy that although no obvious detrimental effect of freezing or delayed sample processing was observed in this study, we did observe some taxonomic changes that can be associated with freezing. For example, in microbiome V31, the relative abundance of genus *Akkermansia* (phylum Verrucomicrobia) in cultured microbiomes was increased when frozen feces were used as inoculum, including those on dry ice for only 6 hours. This might be due to the better resistance of *Akkermansia* species, such as *A. muciniphila*, to stressors including low temperature and oxygen ^32^. Interestingly, Verrucomicrobia has also been frequently reported to be abundant bacteria in microbial communities of sub-ice waters ^33, 34^, indicating their resilience to low temperature and may be more viable following freeze-thaw cycles. Similarly, *Fusobacterium* (phylum Fusobacteria), a type of gram-negative anaerobic bacteria that has been frequently considered as pathogens and being associated with diseases including colorectal cancer ^35-37^, also showed higher relative abundance in cultured microbiome samples using frozen stools as inoculum. These findings indicate the differential impacts of freezing on individual’s microbiome and suggest that the evaluation of biobanked microbiome viability and cultivability is needed for each individual microbiome.

In summary, this study combines an *ex vivo* microbiome assay and metaproteomics to rapidly evaluate the impacts of freezing and delayed sample processing on the cultivability and functional activity of individual microbiome. We showed that the fecal microbiome is stable in deoxygenated buffer with preservant, such as glycerol, on ice for 48 hours, allowing for off-site sample collection and standard two-day shipping to the laboratory. We also demonstrated that freezing the microbiomes at -80°C for one year has minimal detrimental effects on the cultivability of microbiomes and functional responses to treatment with FOS, although changes on some bacterial species, such as those in Verrucomicrobia and Fusobacteria, can be observed in some individual microbiomes. It remains to be studied whether freezing for longer than one year and/or at lower temperature (e.g., liquid nitrogen) affects the functionality of biobanked microbiome. Nevertheless, we recommend that for live microbiota biobanking: 1) fecal samples need to be collected using a deoxygenated buffer with preservant, 2) immediately placed on ice and transferred to the facility within 48 hours, and 3) processed in anaerobic conditions upon reception for biobanking. Our results also suggest that each microbiome needs to be evaluated for its functionality prior to usage for microbiome assay or transplantation.

## Materials and Methods

### Human stool collection, processing, and storage

The protocol for human stool sample collection (# 20160585-01H) was approved by the Ottawa Health Science Network Research Ethics Board at the Ottawa Hospital, Ottawa, Canada. Four healthy volunteers (V31, V33, V34 and V51, with age of 34, 31, 48, and 49 years old, respectively; two men and two women) were recruited for stool sample collection. Exclusion criteria for participation included the presence of irritable bowel syndrome, inflammatory bowel disease, or diabetes diagnosis; antibiotic use or gastroenteritis episode in three months preceding collection; use of pro-/pre-biotic, laxative, or anti-diarrheal drugs in the last month preceding collection; or pregnancy. Fresh stools were collected from the volunteers and were immediately kept on dry ice or immersed in pre-reduced deoxygenated preservation buffer. The deoxygenated preservation buffer was prepared with sterile phosphate-buffered saline (PBS, pH 7.6) with a final concentration of 10% (v/v) glycerol and 0.1% (w/v) L-cysteine hydrochloride. Prior to usage, the preservation buffer was stored in anaerobic workstation (5% H2, 5% CO2, and 90% N2) for overnight with the lid open for gas exchange.

For the first experiment to evaluate the effects of freezing, stools that were kept in deoxygenated preservation buffer were immediately transferred to anaerobic workstation and homogenized to make a 20% (w/v) fecal slurry followed by filtering using sterile gauzes to remove large particles. A minimum of eight aliquots were generated for each microbiome. One aliquot was directly used for RapidAIM culturing (see details below), and the other aliquots were stored in -80 °C for future use at 1, 2, 4, 8, 16, 24, and 52 weeks, respectively. For the second experiment to evaluate the effects of stool transportation on dry ice, stools were kept on dry ice or in deoxygenated preservation buffer on ice for 6 hours. After that, both stool samples were transferred to anaerobic workstation for processing to make a 20% (w/v) fecal slurry as described above. Both samples were then used as inoculum directly for RapidAIM culturing (see details below). For the third experiment to evaluate the effects of delayed sample processing, stools were kept in deoxygenated preservation buffer and stored on ice for 24 or 48 hours prior to sample preprocessing and RapidAIM culturing as described above.

### RapidAIM culturing and metaproteomic sample processing

RapidAIM culturing was performed with a 96-well plate and optimized gut microbiota culture medium as described previously ^11^. The culture medium was composed of 2.0 g L^−1^ peptone water, 2.0 g L^−1^ yeast extract, 0.5 g L^−1^ L-cysteine hydrochloride, 2 mL L^−1^ Tween 80, 5 mg L^−1^ hemin, 5 μL L^−1^ vitamin K1, 1.0 g L^−1^NaCl, 0.4 g L^−1^ K_2_HPO_4_, 0.4 g L^−1^ KH_2_PO_4_, 0.1 g L^−1^ MgSO_4_·7H_2_O, 0.1 g L^−1^ CaCl_2_·2H_2_O, 4.0 g L^−1^NaHCO_3_, 4.0 g L^−1^ porcine gastric mucin, 0.25 g L^−1^ sodium cholate, and 0.25 g L^−1^ sodium chenodeoxycholate. The culture medium was equilibrated in anaerobic workstation overnight before use. Fresh fecal slurry was inoculated into 1 ml microbiota culture medium at a final fecal concentration of 2% (w/v), while frozen fecal slurry was thawed at 37°C with thorough shaking prior to inoculation. FOS was added at a final concentration of 5 mg/ml and PBS (pH 7.6) was added as vehicle control at the same volume of FOS. Following the addition of inoculums, FOS or PBS, the plates were covered with vented sterile silicone mats and shaken at 500 rpm at 37°C for 24 hours in the anaerobic workstation. Baseline samples were collected from the culturing plate after a brief shaking for 2 min after inoculation of microbiota.

Microbial cell harvesting and metaproteomic sample processing were performed according to our previously established workflow ^38^. Briefly, following culturing, samples were transferred from 96-well plates to individual 1.5ml Eppendorf tubes for centrifugation at 14,000 g for 20 min at 4°C to pellet the microbiota cells. The pelleted microbiota cells were then resuspended with 1 mL cold PBS and centrifuged at 300g at 4°C for 5 min to remove any debris. The supernatants were then transferred into new 1.5ml tubes for two additional rounds of debris removal, and the resulting supernatants were centrifuged at 14,000g at 4°C for 20 min to pellet the microbiota cells. The pelleted microbiota cells were washed twice by resuspending in 1 ml cold PBS, centrifugation at 14,000g at 4°C for 20 min followed by supernatant removal. The harvested microbiota cell pellets were then stored at -80°C prior to protein extraction and digestion.

Protein extraction was performed by resuspending the microbiota cell pellets in 100 µL lysis buffer containing 8 M urea, 4% (w/v) sodium dodecyl sulfate (SDS), 50 mM Tris-HCl (pH 8.0), and Roche cOmplete™ Mini protease inhibitor. Lysis was then carried out with three ultrasonications (30 s each with 30 s interval on ice) using QSonica Q125 sonicator (QSonica LLC., Newtown, USA) with an amplitude of 25%. After sonication, samples were centrifuged at 16,000 g at 8 °C for 10 min to remove any cell debris, and the supernatant was transferred to new tube for precipitation using 5 folds volume of ice-cold protein precipitation buffer (acetone/ethanol/acetic acid, 50/50/0.1 (v/v/v)) overnight at -20°C. Proteins were pelleted by centrifugation at 16,000 g at 4°C for 25 min. The proteins were then washed three time by resuspending in ice-cold pure acetone and centrifugation at 16,000 g at 4°C for 10 min with supernatant being removed. After last acetone wash, protein pellets were resuspended in 100 µL of 6M urea in 50mM ammonium bicarbonate for DC^®^ protein assay (Bio-Rad Laboratories) and trypsin digestion.

Fifty micrograms of proteins of each sample were used for in-solution trypsin digestion as described previously ^38^. Briefly, the proteins were first reduced with 10 mM dithiothreitol (DTT) at 56°C for 30 min and alkylated in the dark with 20 mM iodoacetamide (IAA) at room temperature for 45 min. Then the lysates were diluted using 50 mM ammonium bicarbonate to reduce the concentration of urea to less than 1 M. One μg of trypsin (Worthington Biochemical Corp., Lakewood, NJ) and the tryptic digestion was performed at 37°C overnight with agitation. The tryptic peptides were then purified using a 10-μm C18 column and eluted with 80% acetonitrile (v/v)/0.1% formic acid (v/v). After evaporation, the tryptic peptides were dissolved in 0.1% formic acid (v/v) for MS analysis.

### Liquid chromatography-tandem mass spectrometry

Tryptic peptides were analyzed on a Q Exactive mass spectrometer (ThermoFisher Scientific Inc.) coupled with a high-performance liquid chromatography. The separation of peptides was performed on an analytical column (75 μm × 50 cm) packed with reverse phase beads (1.9 μm; 120-Å pore size) with 2-hour gradient from 5 to 35% acetonitrile (v/v) at a flow rate of 200 nl/min. The MS method consisted of one full MS scan from 300 to 1800 m/z followed by data-dependent MS/MS scan of the 12 most intense ions. A dynamic exclusion repeat count was set to 2, and the repeat exclusion duration to 30 s. All data were recorded by Xcalibur version 4.3 and exported into RAW format for further peptide/protein identification and quantification.

Since the experiments in this study span for around one year, the samples were run on MS at different batches. To monitor the performance and consistency of MS measurement, a quality control (QC) sample was prepared by randomly mixing 10 samples and aliquoted for MS run at each batch of MS measurement.

### Bioinformatic data analysis and visualization

All MS raw files were subjected to data processing using MetaLab (version 1.2), a bioinformatic tool for automated and comprehensive metaproteomic data analysis ^39^. Briefly, peptides and proteins were identified and quantified using the MetaPro-IQ workflow ^40^. To generate a reduced database, redundant spectra were removed using a spectral clustering strategy and the resulting clustered spectra were then searched against the human gut microbial gene catalog database (containing 9.9 million microbial genes) ^41^. All matched proteins were then extracted, and their sequences were compiled as a database for a second step target-decoy database search with a strict filtering of the peptide-spectrum matches (PSMs) based on a false discovery rate (FDR) of 0.01. Relative abundances of identified protein groups were quantified using label-free quantification (LFQ) with maxLFQ algorithm ^42^. Taxonomic annotation of all identified peptide sequences was performed using a built-in pep2tax database as described previously ^39^. The identified taxa were then quantified using the intensities of corresponding distinctive peptides and were analyzed at different taxonomic rank levels separately. Identified protein groups were then subjected for functional annotation using eggNOG-mapper ^43^. In this study, Clusters of Orthologous Groups of proteins (COGs) and COG category were used for functional analyses. Relative abundances of a COG or COG category were derived by summing the LFQ intensities of all protein groups that were annotated as that COG or COG category.

Principal component analysis (PCA) was performed in R (version 4.0.2) using the function *prcomp* and visualized using *autoplot* function. To do PCA analysis, quantified protein groups or COGs were first filtered using a criterion of an appearance of non-zero data in ≥ 50% samples; the intensities of the remaining protein groups or COGs were then log_10_-transformed, and the missing values were imputed using K-nearest neighbors algorithm in R with the function *kNN*. Taxonomic compositions at phylum or genus level were visualized using a stacked bar plot with *ggplot2*. Box plots were generated using *ggplot2* as well.

## Data availability

All MS proteomics data along with the MetaLab search results were deposited to the ProteomeXchange Consortium (http://www.proteomexchange.org) via the PRIDE partner repository.

## Acknowledgement

This work was supported by the Government of Canada through Genome Canada and the Ontario Genomics Institute (OGI-156 and OGI-149), the Natural Sciences and Engineering Research Council of Canada (NSERC, grant no. 210034), and the Ontario Ministry of Economic Development and Innovation (ORF-DIG-14405 and project 13440).

